# High Frequency Actionable Pathogenic Exome Mutations in an Average-Risk Cohort

**DOI:** 10.1101/151225

**Authors:** Shannon Rego, Orit Dagan-Rosenfeld, Wenyu Zhou, M. Reza Sailani, Patricia Limcaoco, Elizabeth Colbert, Monika Avina, Jessica Wheeler, Colleen Craig, Denis Salins, Hannes L. Röst, Jessilyn Dunn, Tracey McLaughlin, Lars M. Steinmetz, Jonathan A. Bernstein, Michael P. Snyder

## Abstract

Whole exome sequencing (WES) is increasingly utilized in both clinical and non-clinical settings, but little is known about the utility of WES in healthy individuals. In order to determine the frequency of both medically actionable and non-actionable but medically relevant exome findings in the general population we assessed the exomes of 70 participants who have been extensively characterized over the past several years as part of a longitudinal integrated multi-omics profiling study at Stanford University. We assessed exomes for rare likely pathogenic and pathogenic variants in genes associated with Mendelian disease in the Online Mendelian Inheritance in Man (OMIM) database. We used American College of Medical Genetics (ACMG) guidelines were used for the classification of rare sequence variants, and additionally we assessed pharmacogenetic variants. Twelve out of 70 (17%) participants had medically actionable findings in Mendelian disease genes, including 6 (9%) with mutations in genes not currently included in the ACMG’s list of 59 actionable genes. This number is higher than that reported in previous studies and suggests added benefit from utilizing expanded gene lists and manual curation to assess actionable findings. A total of 60 participants (89%) had non-actionable findings identified including 57 who were found to be mutation carriers for recessive diseases and 21 who have increased Alzheimer’s disease risk due to heterozyg ous or homozygous *APOE* e4 alleles (18 participants had both). These results suggest that exome sequencing may have considerably more utility for health management in the general population than previously thought.

## Introduction

Whole genome sequencing (WGS) and exome sequencing (WES) play increasingly important roles in providing molecular diagnoses for Mendelian disease (Manolio et al. 2013), as well as identifying potential driver mutations in patients with cancer. However, our understanding of the extent to which WGS and WES can benefit healthy individuals is limited. While a few previous studies have attempted to elucidate the utility of WGS in healthy cohorts or individuals (Chen et al. 2012; Xue et al. 2012; Gonzalez-Garay et al. 2013; Dewey et al. 2014; Dewey et al. 2015), more have identified “incidental” or “secondary” findings in disease cohorts—often cohorts with known or suspected genetic disease (Dorschner et al. 2013; Lawrence et al. 2014; Tabor et al. 2014; Amendola et al. 2015; Jang et al. 2015; Jurgens et al. 2015). These studies have reached a wide range of conclusions regarding the rate at which Mendelian disease-causing mutations are identified, due in large part to significant differences in their approaches to variant filtering and curation and the use of gene lists to limit potential findings.

In 2015 the American College of Medical Genetics and Genomics (ACMG) published guidelines to standardize the classification of genomic sequence variants (Richards et al. 2015). These guidelines reinforce the necessity of expert manual curation for accurate variant classification. However, manual curation is labor intensive and has been estimated to take nearly an hour per variant (Dewey et al. 2014). Most previous studies assessing medically relevant WGS and WES findings have classified variants using in-silico predictors or by matching variants against publicly available databases. However, avoiding the step of expert variant curation significantly impairs the ability to accurately classify variants, as in-silico predictors lack accuracy and current publicly available databases for human genomic variants contain variants that are incorrectly classified as disease-causing (Dewey et al. 2014); (Thusberg et al. 2011; Vail et al. 2015; Masica and Karchin 2016). Most previous studies also restricted their analyses by searching for variants in a limited list of genes. However, restricting the search for medically relevant variants to a targeted gene list—for example, the commonly used list of 59 genes compiled by the ACMG—limits findings to only a fraction of genes associated with Mendelian disease (Green et al. 2013; Kalia et al. 2017). Thus, studies that perform a comprehensive analysis of Mendelian risk in generally healthy individuals using ACMG guidelines have not been performed.

In this research study we examine the utility of WES for the general population by using established guidelines to perform an in-depth search for variants with potential medical significance in a group of 70 unrelated adult volunteers enrolled in a longitudinal wellness study. Our analysis included variants in all genes previously associated with Mendelian genetic diseases in the Online Mendelian Inheritance in Man (OMIM) database (http://www.omim.org) or on the list of 59 ACMG genes. In addition, we assessed pharmacogenetic variants. We found a number of medically relevant variants that lie in genes other than those on the ACMG list. These results were reported back to participants by a genetic counselor in accordance with their expressed preferences for the types of results they would like to receive.

## Results

### Participant Demographics

The exomes of 70 participants were analyzed. The participants were all generally healthy at the time of enrollment, with the exception of four diabetics, three of whom were previously diagnosed and being treated, and one with diabetes detected at the time of enrollment due to an HbA1c ≥ 6.5%. Twenty out of 70 participants (29%) were pre-diabetic (defined by a HbA1c between 5.7% and 6.5%), which is similar to the general population prevalence of pre-diabetes (National Diabetes Statistics Report 2014). Participant characteristics are summarized in Table 1. They represented a range of self-reported ethnic backgrounds, including 48 Caucasian, 8 Southeast Asian, 6 Indian, 5 African-American, and 3 Hispanic participants. Thirty-six participants were men and 34 were women. Their ages ranged from 34 to 76 years old with a median age of 57. Fifty-five participants consented to make their sequences public—they are available at http://ihmpdcc.org/resources/osdf.php. These plus the remaining will be made available in dbGAP upon acceptance of the manuscript.

**Table 1:** Participant Demographics

|  | **Range (Median)** |
| --- | --- |
| **Age** | 34-76 (57) |
| **Ethnicity** | **No. of Participants (% of cohort)** |
| Caucasian | 48 (69%) |
| Southeast Asian | 8 (11%) |
| Indian | 6 (9%) |
| African-American | 5 (7%) |
| Hispanic | 3 (4%) |
| **Gender** | **No. of Participants (% of cohort)** |
| Male | 36 (51%) |
| Female | 34 (49%) |

### Exome Results

The gene coding regions were sequenced using an enhanced exome sequencing strategy that provides comprehensive coverage of coding regions as well as additional genomic regions of interest (Patwardhan et al. 2015) (see methods). See methods and Figure 1 for work flow. A range of 149,311 to 262,804 variants was called per exome. Following the filtering steps described in Figure 2, a total of 1,452 variants were reviewed and further filtered manually as described in methods. A total of 668 variants (an average of 9.5 per participant) underwent manual curation using ACMG guidelines (Richards et al. 2015). Of these, 48 variants were classified as pathogenic and 96 as likely pathogenic. The remainder were classified as variants of unknown significance (VUS), likely benign, or benign. The details of variant classification are presented in Table 2.

**Figure 1:**
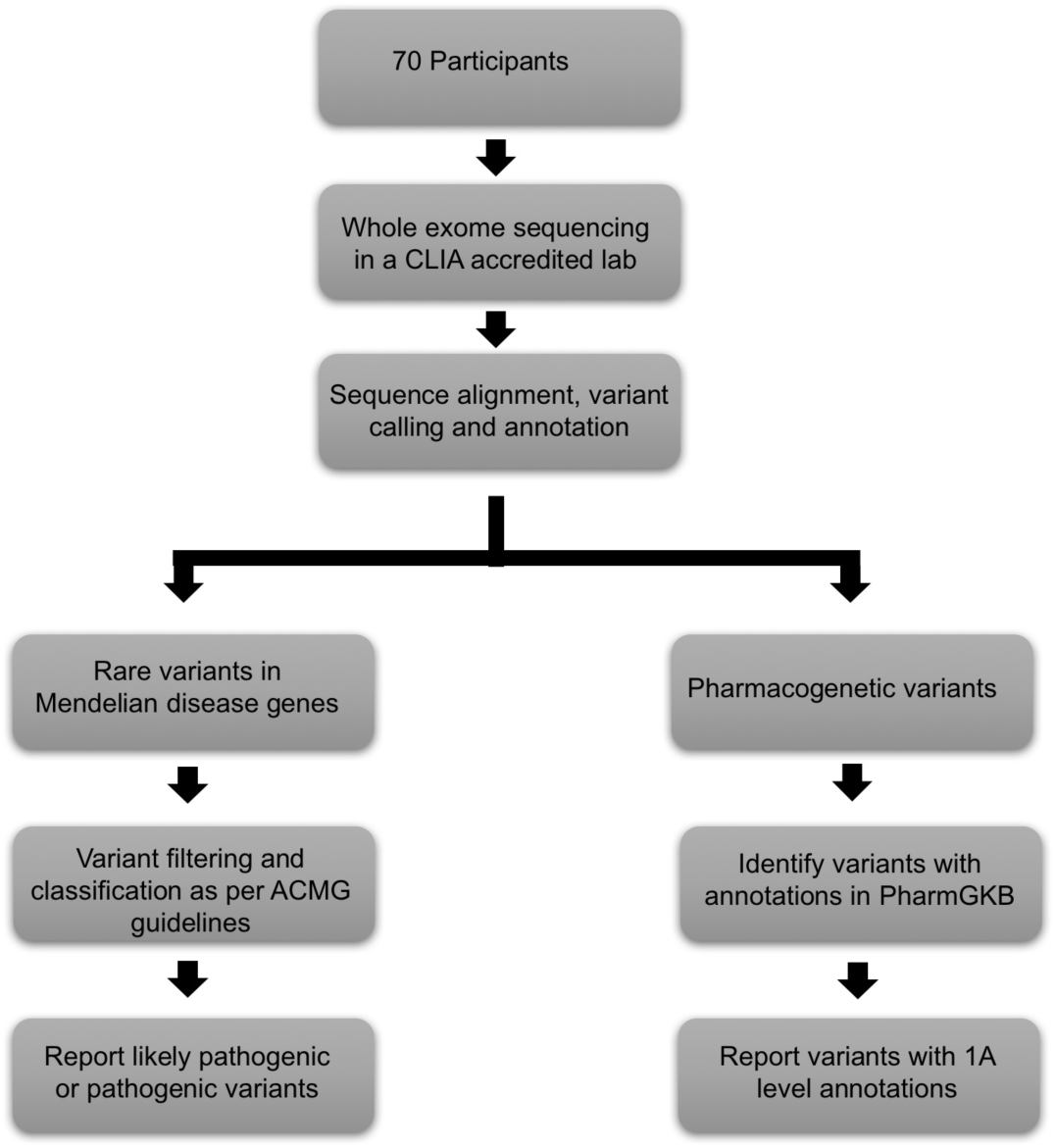
Work Flow

**Figure 2:**
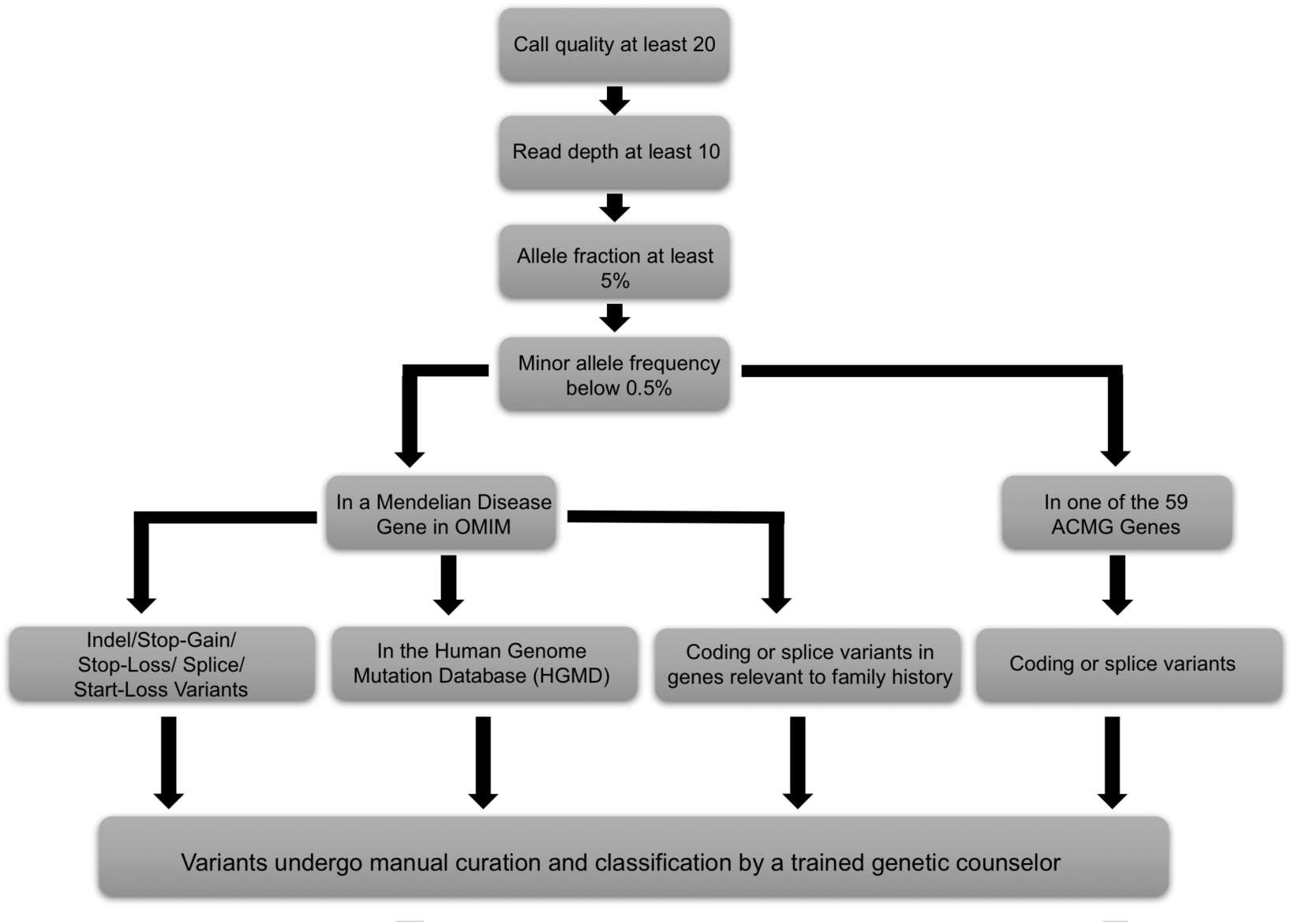
Variant Filtering and Curation

**Table 2:** Variant Classifications

| Variant Call | Number of Variants (average per participant) |
| --- | --- |
| Pathogenic | 48 (0.7) |
| Likely Pathogenic | 96 (1.4) |
| Variant of Unknown Significance (VUS) | 445 (6.6) |
| Likely Benign | 66 (0.9) |
| Benign | 13 (0.2) |
| Reviewed and not classified<sup>a</sup> | 784 (11.2) |

<sup>a</sup>Variants were not classified if viewing the aligned reads suggested the variant was an artifact, if variants in that gene are expected to cause serious, highly penetrant disease at a young age and which the participant did not have the associated phenotype (variants were only removed when the patient had a genotype that would be expected to cause disease were the variant pathogenic—i.e. homozygous for a recessive disease or heterozygous for a dominant disease), or when they were observed in more than 0.5% of a subpopulation in the Exac or 1000 Genomes databases but passed the upstream MAF filter because the overall population MAF was lower than 0.5%

As expected, the vast majority of likely pathogenic and pathogenic variants identified in the cohort were located in genes associated with autosomal recessive diseases, and participants were therefore considered mutation carriers who, in most cases, were unlikely to manifest symptoms. However, actionable pathogenic or likely pathogenic variants were identified 12 participants (see Figure 3). These variants were primarily in genes associated with autosomal dominant disease, although one pathogenic variant was in *MUTYH* (MIM: 604933)—a gene which is associated with autosomal recessive *MUTYH*-associated polyposis (MIM: 608456)—but for which carriers are known to be at increased lifetime colon cancer risk (5.6% for female heterozygotes and 7.2% for male heterozygotes by age 70; higher for patients with a first degree relative with colon cancer) (Win et al. 2014). Due to this increased risk, the National Comprehensive Cancer Network (NCCN) has issued screening guidelines for heterozygous mutation carriers (National Comprehensive Cancer Network 2016). Therefore, we considered this variant actionable. The actionable variants lie in 10 distinct genes (Table 3) and include five variants classified as pathogenic with strong evidence suggestive of a causative role in disease as per ACMG classification guidelines; five classified as likely pathogenic; and one variant that was identified in two individuals was classified as a risk allele. The risk allele—in the *APC* gene (MIM: 611731)—is a well-studied founder mutation in the Ashkenazi Jewish population that the NCCN has described as a moderate risk allele for colon cancer, and has issued screening guidelines for heterozygous carriers of this mutation (Boursi et al. 2013; Liang et al. 2013; National Comprehensive Cancer Network 2016). In total, 12 of the 70 individuals in the cohort (17%) had medically actionable likely pathogenic or pathogenic variants identified (see Table 3 for the complete list of actionable variants). Of the 12 variants, six reside in the 59 genes reported as actionable in the most recent ACMG guidelines regarding incidental findings (Green et al. 2013; Kalia et al. 2017). These include heterozygotes for likely pathogenic and pathogenic mutations in the highly penetrant cancer risk genes *BRCA1* (MIM:113705), which is associated with hereditary breast and ovarian cancer (MIM: 604370) and *SDHB* (MIM: 185470), which is associated with hereditary paraganglioma and pheochromoctytoma (MIMs: 115310, 171300). The remaining six variants reside in genes that are not included in the ACMG guidelines but that are associated with medically actionable disease as defined in the methods.

**Figure 3:**
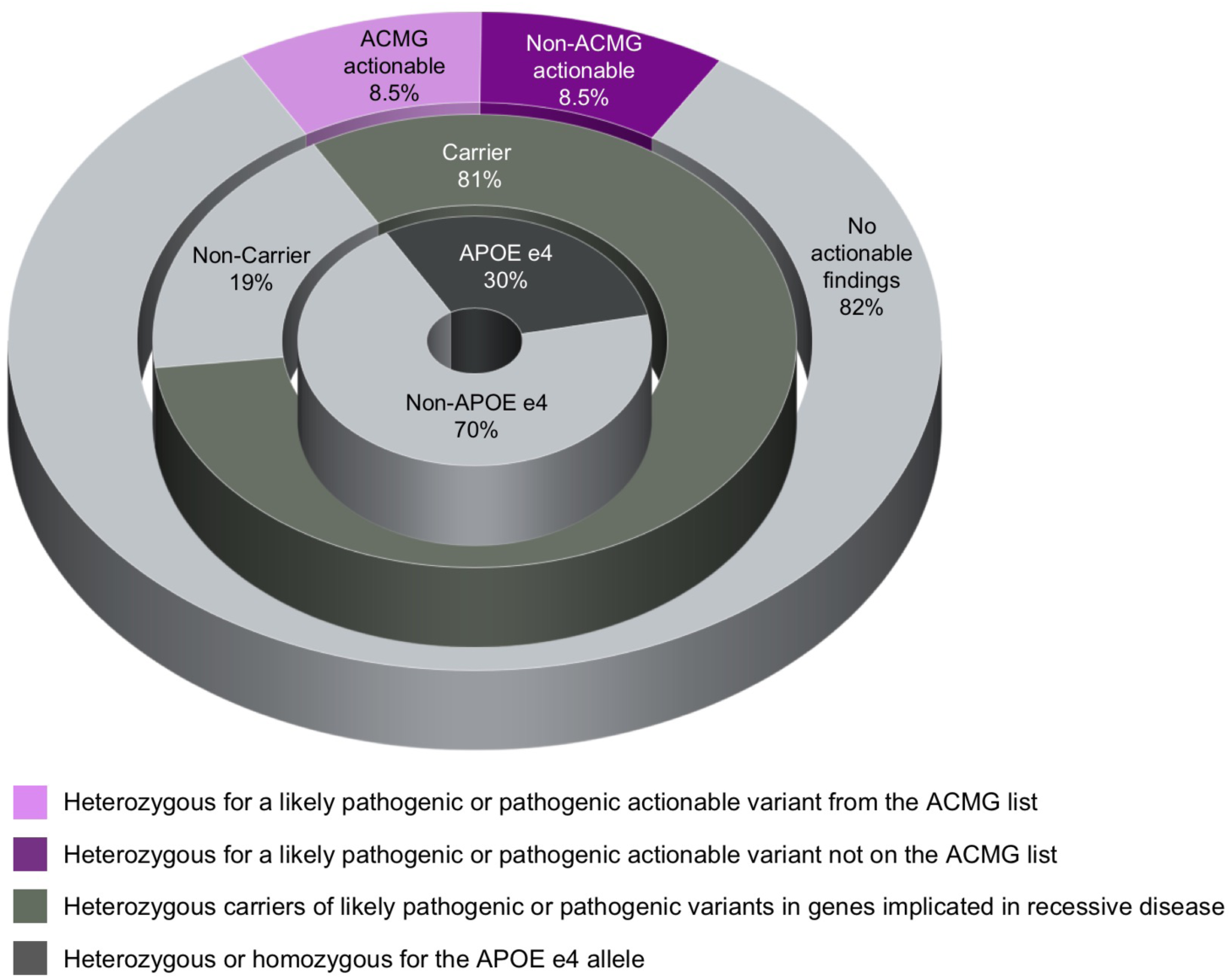
Actionable and non-actionable exome findings

**Table 3:**
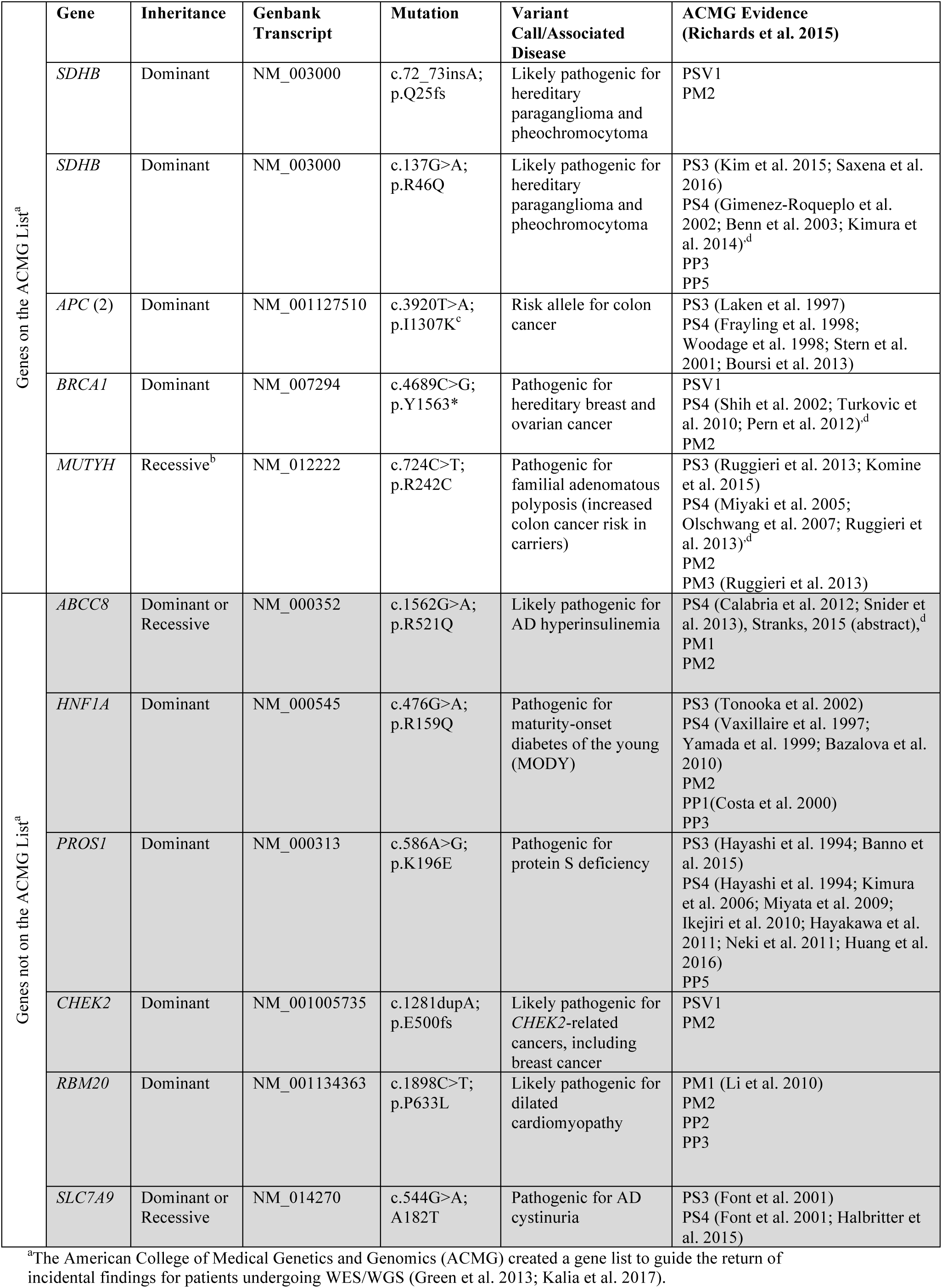
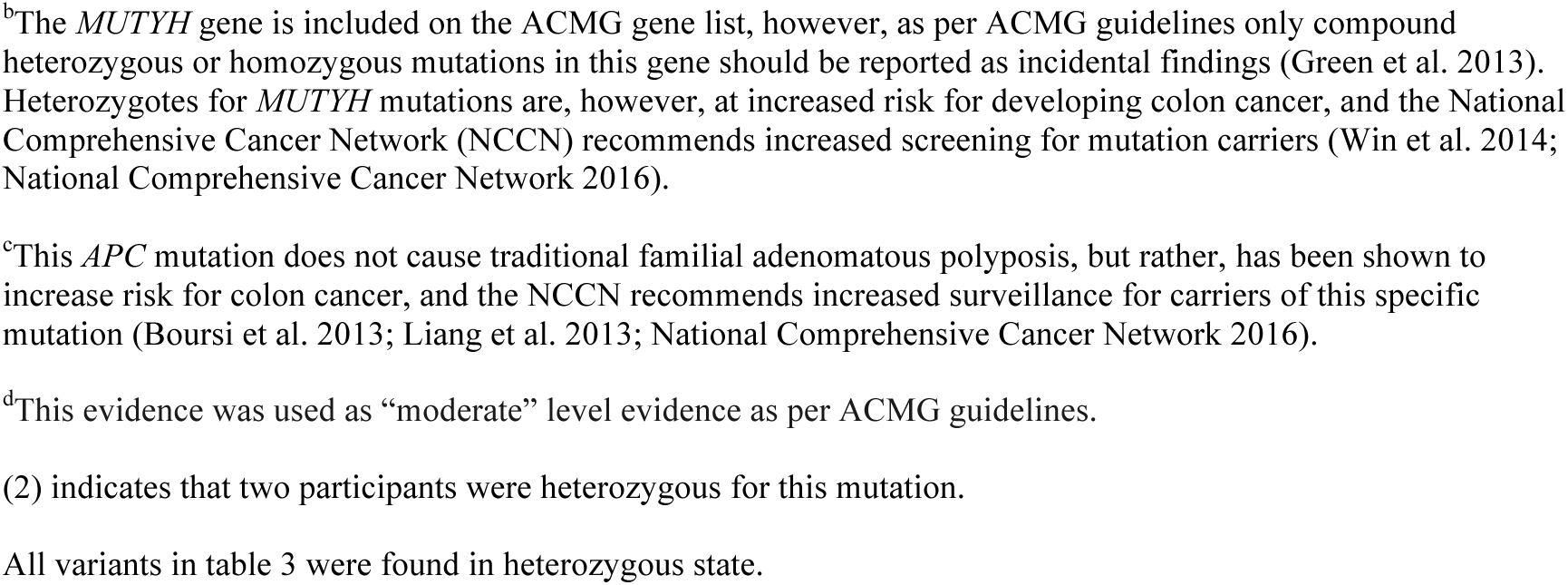
Medically Actionable Exome Findings

### Patients with Personal or Family Medical History Consistent with their Mutation

At least two individuals have personal or family medical histories consistent with the presence of their mutations. A 46-year-old female with elevated glucose and family history of diabetes was found to be heterozygous for a likely pathogenic *HNF1A* (MIM: 142410) mutation. *HNF1A* mutations cause autosomal dominant maturity onset diabetes of the young (MODY [MIM: 600496]), a form of monogenic diabetes that is often misdiagnosed as type 1 or type 2 diabetes, as was the case in this participant, who was incorrectly diagnosed with type 2 diabetes in her early 30s. Compared to type 2 diabetes, diabetes caused by *HNF1A* mutations is considerably more responsive to sulphonylurea drugs. Early recognition of *HNF1A*-MODY and subsequent initiation of sulphonylurea therapy may reduce the incidence of diabetic complications (Shepherd et al. 2009; Bacon et al. 2016). The participant, who was currently managing her diabetes with a combination of three non-sulphonylurea oral medications, was referred to endocrinology to discuss potential changes to her treatment plan, and her three children also underwent genetic testing for the mutation to inform diabetes screening regimens.

Another participant with a family history of dilated cardiomyopathy (DCM [MIM: 613172]) was identified to be heterozygous for a likely pathogenic *RMB20* (MIM: 613171) variant. The variant has not been previously reported as associated with DCM but was prioritized for curation as a result of the participant’s family history. The variant is in a hotspot for DCM-associated mutations located in the RS domain of *RBM20* and is located in a codon adjacent to a series of five codons previously reported in DCM cases (Li et al. 2010). Due to family history, as well as low-normal ejection fraction on echocardiogram revealed due to follow up subsequent to genomic analysis, the participant began taking blood pressure lowering medications as a preventative measure. The participant was referred to cardiovascular genetics clinic for follow up.

### Non-medically actionable findings

A total of 60 participants (89% of the cohort) were identified to have non-actionable findings (including carriers for recessive conditions and/or *APOE* e4 allele carriers—see Figure 3). In 57 participants we identified 133 likely pathogenic and pathogenic variants in genes that cause autosomal recessive diseases (see supplemental table 1 for a complete list of likely pathogenic and pathogenic variants). Most of these variants convey no health risks to carriers beyond reproductive risks, but there are exceptions. In addition to the *MUTYH* mutation discussed earlier, pathogenic heterozygous *GBA* (MIM: 606463) mutations were identified in three participants. *GBA* mutations cause autosomal recessive Gaucher disease (MIMs: 608013, 230800, 230900, 231000, 231005), but like individuals affected with Gaucher disease, heterozygous mutation carriers are also at significantly increased risk for Parkinson’s disease (MIM: 168600)(Tayebi et al. 2003; Halperin et al. 2006; Alcalay et al. 2014). In addition, 21 participants were identified to be heterozygous or homozygous for the *APOE* (MIM: 107741) e4 allele, which is associated with significantly increased lifetime risk for developing Alzheimer’s disease (MIM: 104310) (Corder et al. 1993; Bertram et al. 2010). The *APOE* genotype was only disclosed in two cases where participants specifically inquired about it and had opted to receive both actionable and non-actionable findings on their consent form.

### Pharmacogenetic variants

In addition to disease causing variants we also assessed participant exomes for variants impacting response to drugs. Level 1A variants in PharmGKB (https://www.pharmgkb.org) have high confidence for affecting drug dose and/or side effects. The 70 exomes were examined for 28 rsIDs with level 1A classifications (extracted from pharmgkb.org in May 2017) (see supplemental table 2 for a list of rsIDs). A range of 1 to 6 level 1A variants were identified per participant, with a median of 3 variants. Well-known examples include several variants in *CYP2C19* that are associated with altered metabolism or risk of side effects for drugs such as Clopidogrel and Amitriptyline, including rs9923231 and rs4244285 (Whirl-Carrillo et al. 2012). Thus, overall, the majority of our participants received potentially useful pharmacogenetic information.

## Discussion

Previous studies have found that approximately 1-5% of individuals will have actionable incidental or secondary findings identified on exome sequencing (Johnston et al. 2012; Dorschner et al. 2013; Lawrence et al. 2014; Amendola et al. 2015; Jurgens et al. 2015). The substantially higher rate of actionable findings found in our cohort (17%) suggests that employing manual variant curation and expanded gene lists may enhance the accurate identification of actionable variants in exomes—information that can ultimately lead to improved treatment, screening, or prevention. Our study also highlights the fact that many individuals can receive useful risk information from genome and or exome sequencing.

### The ACMG List and Beyond

A number of previous studies of the utility of WGS or WES in healthy individuals have limited their search for clinically relevant findings to the list of 56 genes first compiled by the ACMG in 2013 and later revised to include 59 genes in 2016 (Green et al. 2013; Kalia et al. 2017). It is recommended that incidental findings from genes on this list be returned to patients regardless of the indication for WGS/WES, as pathogenic/likely pathogenic variants in these highly penetrant genes lead to high risk for serious preventable and/or treatable diseases. The wide range of percentages (1-5%) of actionable findings reported in previous studies may in part be explained by the lack of widely accepted standardized guidelines for interpretation of genomic variants prior to 2015 (Richards et al. 2015). Although variants in the 59 ACMG genes are actionable and therefore clinically relevant, they represent only a fraction of genes known to cause Mendelian disease in humans.

Among the criticisms of the ACMG list is the lack of agreement among experts on the ACMG panel that compiled the list over which findings should be reported, as well as the existence of numerous other genes in which mutations can cause highly penetrant and treatable or preventable disease (Burke et al. 2013; Holtzman 2013). Our findings, which include several medically actionable pathogenic or likely pathogenic variants in genes not included in the latest version of the ACMG list, support this concern. In addition to mutations in genes on the ACMG list we identified pathogenic or likely pathogenic variants in six other genes not on the ACMG list that have medical relevance, including two (*HNF1A* and *RBM20*) in which participants had personal and/or family medical history consistent with pathogenicity.

In addition to *HNF1A* and *RBM20* mutations, we identified mutations in other noteworthy genes. As previously noted, one participant was found to carry a likely pathogenic *MUTYH* mutation. Current ACMG guidelines for the reporting of incidental findings recommend only reporting compound heterozygote or homozygous mutations in *MUTYH*, as *MUTYH-* associated polyposis is considered a recessive disease (Green et al. 2013; Kalia et al. 2017). However, heterozygote carriers of *MUTYH* mutations are at an increased risk of colon cancer and the National Comprehensive Cancer Network (NCCN) has recommended carriers undergo more frequent colonoscopies starting at an earlier age than the general population recommendations (National Comprehensive Cancer Network 2016). In one male participant we identified a pathogenic *CHEK2* mutation, which leads to a dominantly-inherited high lifetime risk for cancers including breast and colorectal (Meijers-Heijboer et al. 2002; Xiang et al. 2011). *CHEK2* is not on the ACMG list, but NCCN recommends increased cancer screening starting at younger ages for mutation carriers (National Comprehensive Cancer Network 2017). Identification of such mutations can also alert family members to their potential cancer risk, which for female relatives of our participant found to carry his same mutation would include a significant (potentially more than two-fold) increased breast cancer risk (CHEK2 Breast Cancer Case-Control Consortium 2004). Both of these participants were referred to a cancer genetics clinic for follow up.

Additionally, a female participant found to be heterozygous for a pathogenic *PROS1* mutation is at increased risk for thrombophilia due to protein S deficiency, which leads to preventative treatment in some patients, particularly those who already have a family history of thrombotic events (De Stefano and Rossi 2013). Oral contraceptives are also contraindicated in women with heterozygous *PROS1* mutations, even in the absence of family history of thrombotic events (van Vlijmen et al. 2016).

While the ACMG list has become the default gene list used to determine which incidental/secondary findings should be returned to participants undergoing WES/WGS, it is not necessarily a comprehensive representation of all such genes. Additionally, a number of studies have demonstrated that many patients and research participants do want to learn about incidental or secondary findings that are not medically actionable—such as genetic risk for developing adult-onset neurodegenerative conditions such as Alzheimer’s disease or Parkinson’s disease. Qualitative research on this subject has suggested many individuals find this information actionable in other (non-medical) ways and express that they would live their lives differently if they knew they were at increased risk of developing such a condition or would prepare for developing the disease (Clift et al. 2015; Yushak et al. 2016). Our study identified 21 participants with one or two copies of the *APOE* e4 allele, which significantly increases lifetime risk for developing Alzheimer’s disease (Corder et al. 1993; Bertram et al. 2010). This information was reported back to participants who specifically requested their *APOE* status. Similarly, we identified three heterozygous carriers of *GBA* mutations, and while *GBA* carriers will not develop Gaucher disease—an autosomal recessive lysosomal storage disease—they are at increased risk for developing Parkinson’s disease (Tayebi et al. 2003; Halperin et al. 2006; Alcalay et al. 2014). We reported this information back to participants who opted to learn all medically relevant findings. For the *GBA* mutation carriers in our study as well as the 54 carriers of mutations in other genes implicated in recessive disease, this information can also alert families to potential reproductive risk and lead to carrier testing for their partners or adult children.

### Manual Variant Curation

In addition to limiting incidental findings to variants within a specific gene list, researchers attempting to look for WGS/WES secondary findings have also attempted to mitigate the curation workload by forgoing variant curation altogether. Several previous studies assessing WGS/WES secondary findings have either completely or primarily relied on a combination of in-silico predictors and variant databases such as Clinvar (https://www.ncbi.nlm.nih.gov.clinvar) and HGMD (www.hgmd.cf.ac.uk/ac/index.php) to classify variants rather than employing manual curation (Gonzalez-Garay et al. 2013; Tabor et al. 2014; Gambin et al. 2015; Dewey et al. 2016). This approach may be limiting and error-prone. Testing well-known, previously classified missense mutations with the commonly used in-silico predictors SIFT and Polyphen yields accuracy ranging from 62-78% (Masica and Karchin 2016). Splice site predictors are only slightly more accurate (Vreeswijk et al. 2009; Houdayer et al. 2012). Although improving, the majority of variants in Clinvar have not undergone expert review and classifications are often based on incomplete or outdated evidence and/or were classified without applying stringent criteria. Similarly, variants listed as disease mutations (DM) in the HGMD frequently do not meet criteria to be classified as likely pathogenic or pathogenic. Dewey et al found that only one-fourth of the HGMD DM variants they identified in their cohort were classified by experienced curators as likely pathogenic or pathogenic (Dewey et al. 2014). This evidence supports the necessity of manual variant curation in order to accurately classify variants, at least until such a time as a reliable publicly available database exists.

### Limitations

Our study had several limitations. Among them, we used a minor allele frequency cutoff of 0.5% when filtering variants for further curation, and this will certainly lead to the exclusion of some particularly common pathogenic and likely pathogenic variants, such as the common delta 508 *CFTR* mutation. Other filtering cutoffs also likely limited the number of disease-causing mutations identified. Our cohort size (70) is small and larger studies will be needed to determine if the rate of actionable findings identified in our cohort also applies to larger populations. Our understanding of penetrance in many disease genes is based largely on studies of families known to be affected with disease, so in the future we may learn that penetrance is lower for individuals without family histories of disease who have actionable mutations described in this study.

### Conclusions

We demonstrate that exome sequencing of participants in a longitudinal wellness study reveals important medical information in a considerable fraction of the population. The exome results from our cohort include several medically actionable variants in genes not included in the ACMG list, which suggests a need for a more comprehensive list of genes to guide the return of incidental findings. Indeed, an expanded list of genes for the return of incidental findings would present challenges, as a larger list would require more resources to accurately curate and classify variants and could lead to costly follow-up. More research is needed to better understand how common medically actionable variants are, which other types of results patients find utility in, and to better understand the costs and benefits of returning more extensive secondary findings to patients undergoing exome or genome sequencing. Nonetheless, our study provides a general approach for how to use genome sequencing, interpretation and reporting of medically relevant variants for the population at large.

## Methods

### Recruitment and Study Population

Participants were enrolled as part of Stanford’s iPOP (Integrated Personal Omics Profiling) research study (IRB 23602), which entails longitudinal multi-omics profiling of a cohort of unrelated adult volunteers enriched for pre-diabetics. The iPOP study has been described previously (Chen et al. 2012; The Integrative Human Microbiome Project 2014). All research participants received genetic counseling by a medical geneticist or genetic counselor prior to enrollment and signed a consent form approved by the Stanford University Institutional Review Board. Participants were able to opt in or out of receiving incidental findings, and if they opted in, were also given the option of selecting whether they wanted only actionable results or all results with medical relevance.

### Whole Exome Sequencing

Whole exome sequencing was performed on 70 individuals. Briefly, DNA was isolated from blood using Gentra Puregene Kits (Qiagen) according to manufacturer’s protocol. Exome sequencing was performed at Personalis—a CLIA- and CAP-accredited facility—using the ACE Clinical Exome Test which covers exomes in a more comprehensive fashion (Patwardhan et al. 2015) and additional genomic regions of interest. Variants were called using an updated version of the HugeSeq pipeline (Lam et al. 2012) which used GATK3.3 or higher (McKenna et al. 2010).

### Variant Filtering and Analysis

The overall workflow is depicted in Figure 1. Two types of genomic results were assessed— rare variants in known Mendelian disease genes and variants with pharmacogenetic annotations in the PharmGKB database (https://www.pharmgkb.org). Rare variants were filtered according to the steps depicted in Figure 2. Initially variants were filtered based on confidence metrics including Phred scores (minimum 20) and read depth (minimum 10). Variants were also excluded if they had a minor allele frequency higher than 0.5% in the 1000 Genomes database (www.internationalgenome.org/1000-genomes-browsers) or Exome Aggregation Consortium (ExAC) database (www.exac.broadinstitute.org). We then removed variants that did not appear in one the 3,651 genes in the OMIM database categorized as a gene associated with Mendelian disease (downloaded January 2016), or on the list of 59 genes in which the ACMG recommends reporting incidental findings (Green et al. 2013; Kalia et al. 2017). Additional filtering was performed by 1) the removal of variants for which manual examination of the aligned reads indicated a likely sequencing error; 2) the removal of variants expected to cause serious, highly penetrant disease at a young age and for which the participant did not have the associated phenotype (variants were only removed when the patient had a genotype that would be expected to cause disease were the variant pathogenic— i.e. homozygous for a recessive disease or heterozygous for a dominant disease); our experience revealed that these were usually artifacts; 3) when the curators determined there was insufficient evidence that the gene in which the variant resided was associated with disease; and 4) when the minor allele frequency of the variant in the 1000 Genomes or ExAC database was above 0.5% in a subpopulation but had initially passed filtering because the overall population minor allele frequency was below that cutoff. The remaining rare variants then underwent manual curation and classification by a trained genetic counselor according to ACMG criteria for the classification of sequence variants: 1) variants of a type likely to cause loss of gene function (insertions and deletions, nonsense, splice), 2) variants with an exact match in the Human Genome Mutation Database (HGMD), and 3) coding or canonical splice-site variants in one of the 59 genes in which ACMG recommends reporting incidental findings (Richards et al. 2015; Kalia et al. 2017).

Participants had varying degrees of personal and family medical history available for the curators to take into consideration when classifying variants. For some participants this information was limited to a medical history intake form and basic medical records; for others much more extensive medical history and/or a three-generation pedigree were available. Additional variants were curated when they were identified in genes associated with a potentially Mendelian disease in the participant’s family history.

Participants in whom medically significant likely pathogenic or pathogenic variants were identified were encouraged to discuss the results with their physician and, when necessary, referred to a genetics clinic for follow up. Participants were given the option at the time of consent of selecting whether they would like to receive genomic results, and if so, whether they would prefer actionable results only or all medically relevant results identified. Actionable results were defined as likely pathogenic or pathogenic variants in genes associated with diseases that are moderately to highly penetrant, the identification of which was likely to result in altered medical care in the form of treatment, screening, or preventative measures. Additionally, non-actionable findings with medical relevance were returned to participants who opted to receive them during the consent process. These results included likely pathogenic and pathogenic heterozygous variants in genes implicated in recessive diseases, as well as likely pathogenic and pathogenic variants in genes associated with diseases such as Parkinson’s disease or Alzheimer’s disease, for which limited or no highly effective treatment or preventative measures are available. All pathogenic and likely pathogenic variants were reviewed by two genetic counselors and a medical geneticist. Results were then reported back to participants by a genetic counselor in accordance with their stated preferences.

Participants’ genotypes were also examined for common SNPs with pharmacogenetic annotations that reached a level 1A classification in the PharmGKB database (pharmgkb.org). Level 1A variants represent those with the highest level of validation.

## Data Access

Data from participants who consented to make their sequences completely public is available at http://ihmpdcc.org/resources/osdf.php. Data will also be submitted to dbGAP prior to publication.

## Acknowledgments

Our work was supported by grants from the NIH Common Fund Human Microbiome Project (HMP) (1U54DE02378901) (M.P.S. and T.M.) and American Diabetes Association (grants 1-14-TS-28 and 1-11-CT-35) (T.M.). M.R.S and H.L.R. are supported by grants from the Swiss National Science Foundation (SNSF: P300PA_161005, P2GEP3_151825, M.R.S.; P300PA_164703, H.L.R.). J.D. is funded by the Mobilize Center (grant NIH U54 EB020405). This work was also supported by a gift from the Forbes Family Fund. The authors would like to thank the Stanford Genetics Bioinformatics Service Center for computational and informatics support, as well as the volunteers who participated in our study

